# In silico self-assembly and complexation dynamics of cationic lipids with DNA nanocages to enhance lipofection

**DOI:** 10.1101/2025.06.11.659007

**Authors:** Sandip Mandal, Dhiraj Bhatia, Prabal K. Maiti

## Abstract

DNA nanostructures are promising materials for drug delivery due to their unique topology, shape, size control, biocompatibility, structural stability and blood-brain-barrier (BBB) penetration capability. However, their cellular permeability is hindered by strong electrostatic repulsion from negatively charged cellular membranes, posing a significant obstacle to the use of DNA nanostructures as a drug delivery vehicle. Recent experimental studies have shown enhanced cellular uptake for the conjugate binary mixtures of DNA Tetrahedron (TDN) with cationic lipid N-[1-(2,3-dioleyloxy)propyl]-N,N,N-trimethylammonium chloride (DOTMA) compared to TDN alone. However, the cationic DOTMA lipids binding mechanism with the TDN nucleotides is still elusive. Using fully atomistic MD simulations, we aim to understand the molecular interactions that drive the formation and stability of the TDN-DOTMA binary complexes in a physiological environment. Our results uncovered that lipid concentration plays a crucial role in the energetics of the TDN-DOTMA association. We also report that distinct timescales are associated with the self-assembly of cationic DOTMA lipids first, followed by the complexation of self-assembled DOTMA lipid clusters with the TDN nucleotides, where electrostatics, hydrophobicity, and hydrogen bonding are the key interactions that drive the formation and stability of these complexes. These results provide molecular insights into TDN-DOTMA interactions, highlighting the lipid self-assembly dynamics, complex stability, and morphology, paving the way for the better rational design of cationic lipid functionalized DNA nanostructures for efficient drug delivery and transfection.

## I. INTRODUCTION

DNA nanotechnology, a field pioneered by Prof. Ned Seeman, allows the creation of nanostructures with precise control and programmability of various shapes and sizes via Watson-Crick base-pairing (A-T and G-C). The field has optimized the synthesis of various geometries such as tetrahedral, cubic, icosahedral, bucky ball, and nanotubular shapes, demonstrating the versatility of programmable DNA self-assembly^1–8^. These DNA nanocages have gained popularity in biological and biomedical applications due to their structural stability, programmability, biocompatibility, biodegradability, low cytotoxicity, and unique topology. Their characteristics, such as high payload capacity and customizable designs for targeted release, make them ideal carriers for drug delivery, cellular biosensing, and other therapeutic applications^9,10^.

DNA Tetrahedron (TDN), one of the DNA nanocages, has shown promising abilities as a vehicle for efficient drug delivery due to its ability to encapsulate cargo, penetrate cells, and release payloads in a controlled manner^10–12^. It can enter the cells by crossing the cellular barrier without the use of a transfection agent. TDN was also observed to be a geometry preferred by the cells among an array of four different geometries, as shown by Rajwar et. al.^13^. However, a major challenge in using DNA-based TDN lies in its strong negative charge, which might lead to repulsion from the similarly charged cell membrane. It might be a hindrance while transporting drugs in its hydrophobic core. One way to overcome this major limitation is by functionalizing TDN with a positively charged molecule.

Lipids are an integral part of cellular plasma membranes, accounting for 50% of the total mass of cellular membranes. Cationic lipids, which contain a positively charged head group, help in increasing the efficiency of different materials entering the cell^14–16^. Non-viral vectors such as DOTMA are seen as more promising delivery agents, and DOTMA can help in increasing the uptake efficiency of TDN. However, using cationic lipids as effective transfection vectors with TDN is associated with challenges, particularly the instability of lipid/DNA complexes in biological environments, as previously reported^17,18^. These challenges are influenced by the structural and dynamic properties of lipids, as well as their interactions with DNA^19^. Therefore, gaining mechanistic insights into their self-assembly and complexation will offer a framework for optimizing lipid-functionalized DNA nanostructures for more efficient drug delivery platforms.

A complete knowledge of the nature of atomic-level interactions within TDN-DOTMA complexes remains elusive due to the limitations of current experimental techniques, despite significant progress in both in vivo and in vitro studies^20^. No atomistic MD study involving DOTMA with or without DNA is reported. Only a few studies have attempted a full description of DNA nanostructures in the all-atom form due to the inherent challenges associated with the models’ complexity and the scarce availability of powerful computer resources. In this work, we studied the associative behavior between cationic lipid DOTMA and TDN in an aqueous environment at physiological salt concentrations using atomistic MD simulations. Our primary objectives were to gather comprehensive information on the key steps of the TDN-DOTMA and DOTMA-DOTMA complexation process and assess their potential as nucleic acid delivery platforms and components of LNPs.

## II. MODELING AND METHODOLOGY

### A. System setup

The initial atomic coordinates of the DOTMA lipid structure were obtained from the PubChem database, corresponding to the PubChem CID: 6913074. The geometry optimization of the DOTMA molecule was then carried out using the B3LYP functional together with 6-311G* basis set in the Gaussian 09 program. RESP charges were calculated for these geometry-optimized DOTMA structures (Figure S.1 in the supplementary material). The initial atomistic structure of the TDN cage was built using the Polygon code^21^, and the corresponding base sequences were obtained from the earlier experimental work^20^.

We have investigated five different systems, starting with an aqueous solution of 10 and 100 DOTMA lipid molecules, which were used as the reference system. The other three systems were comprised of 1, 10, and 100 DOTMA lipids along with TDN nanostructure, water, and 150 mM of MgCl_2_ ions in the simulation box as listed in Table I. The built structures were then solvated in a cubic box with a water thickness of 15 Å from the solute using the TIP3P^22,23^ water model using the xLEAP module of AMBER20^24–26^. In this way, all the atoms of the solute molecules will be 15 Å away from the edge of the water box, ensuring that the molecule does not interact with its periodic images. The negative charges of the phosphate backbones were neutralized by adding the appropriate number of Mg^2+^ counter ions. To achieve systems with the desired physiological salt concentration of 150 mM, we added appropriate numbers of MgCl_2_ ions. Periodic boundary conditions were maintained in all three dimensions to mimic the bulk properties of the system. All the MD simulations were carried out using PMEMD.cuda module of the AMBER20 software. The interactions of TDN atoms along with TIP3P waters are described by the AMBER bsc1^27^ forcefield. For the divalent (Mg^2+^) ions, the Li-Merz ion parameters were chosen^28^. The interaction parameters for the DOTMA atoms were generated from the general Amber forcefield (GAFF^29^) with the restrained electrostatic potential (RESP) charges^30^.

### B. Simulation protocols

The energies of the solvated systems were first minimized using the steepest descent algorithm for the first 5000 steps, followed by the conjugate gradient algorithm for another 5000 steps. During the energy minimization, all of the solute atoms were restrained by a harmonic potential with a force constant of 1000 kcal.mol^−1^.Å^−2^ so that ions and water molecules can redistribute themselves to remove any kind of bad contacts.

The systems were subject to gradual heating in four steps: first, from 10 K to 50 K for 6000 steps, then from 50 K to 100 K for 12,000 steps, then from 100 K to 200 K for 10,000 steps, and finally, from 200 K to 300 K for 10,000 steps. Positional restraints were applied to the heavy atoms of the solute during heating with a force constant of 50 kcal.mol^−1^. Å^−2^. Following the heating, systems were subject to a 5 ns NPT equilibration with a Berendsen weak coupling method^31,32^ to adjust the bulk water density with a target pressure of 1 bar. All bonds involving hydrogens were constrained using the Shake algorithm^33^. This allowed us to use an integration time step of 2 fs. The long-range electrostatic interactions were handled using the PME^34^ scheme with a cutoff of 10 Å. The short-range electrostatics and vdW interactions were captured with a distance cutoff of 10 Å. Following the equilibration, the systems were simulated in the NVT ensemble using a Langevin thermostat^32,35^ with a coupling constant of 1 ps at 300 K, for the production run of 200 ns. Similar simulation methodologies at ambient temperatures have been implemented in some of our previous studies^10,36–40^.

All the MD simulation trajectories were visualized with the VMD^41^, CPPTRAJ^42^ for modeling and the analysis of molecular systems. Pymol and Python Matplotlib^43^ modules were also used to visualize and plot the simulation data and the construction of the figures.

## III. RESULTS AND DISCUSSION

### A. Structural Properties

A clear understanding of cationic lipid organization and complexation is crucial for designing lipid-modified drug delivery systems like TDN for nucleic acid-based therapeutics. Here, we focus on the morphology of self-assembled structures formed by DOTMA lipids alone and in complexation with TDN. Due to their hydrophobic nature, DOTMA lipids readily self-assemble in aqueous environments and are also attracted towards negatively charged TDN nucleotides. Therefore, we aim to investigate: (i) how TDN’s negative charge affects DOTMA-DOTMA self-assembly, (ii) whether the complexation of cationic lipids with TDN reduces its negative electrostatic potential, which will enhance its gene delivery capability, and (iii) the key factors that contributes to the stable complex formation of TDN-DOTMA.

Figure 1A-C shows how TDN cage associates with 1, 10, and 100 DOTMA lipids, respectively. Our results indicate that a single DOTMA lipid, initially placed randomly in the simulation box, binds to the outside corner mode of TDN and forms a stable TDN-1 DOTMA complex as shown in Figure 1A. The outside corner mode of TDN is also a prominent binding location as reported previously by Xu et al.^11,12^. For higher DOTMA ratios of 10 and 100 lipids with single TDN cage, we observe that lipids first form self-assembled clusters, which then associate with TDN nucleotides, unlike the interaction of individual DOTMA lipids with the oppositely charged TDN nanostructures. To compare the effect of the presence of TDN in the DOTMA self-assembly, we have also simulated two systems with 10 and 100 cationic DOTMA lipids in bulk water. As shown in Figure 2A–B, both formed stable clusters within the first 50 ns.

**FIG. 1.**
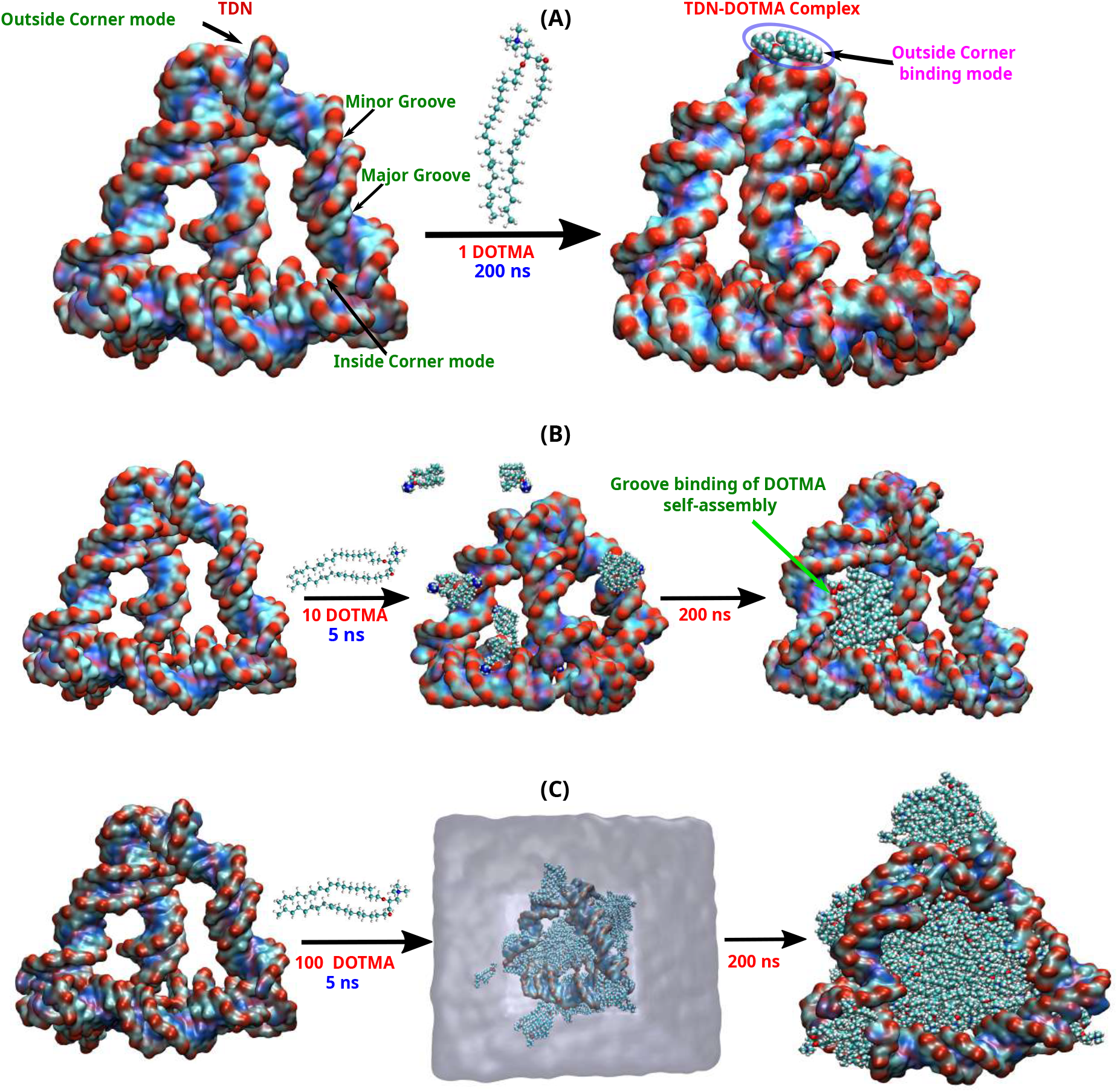
Instantaneous snapshots of the initial and final frames from 200 ns atomistic MD simulations of TDN-DOTMA complexes are shown for systems with (A) 1, (B) 10, and (C) 100 DOTMA lipids, respectively.

**FIG. 2.**
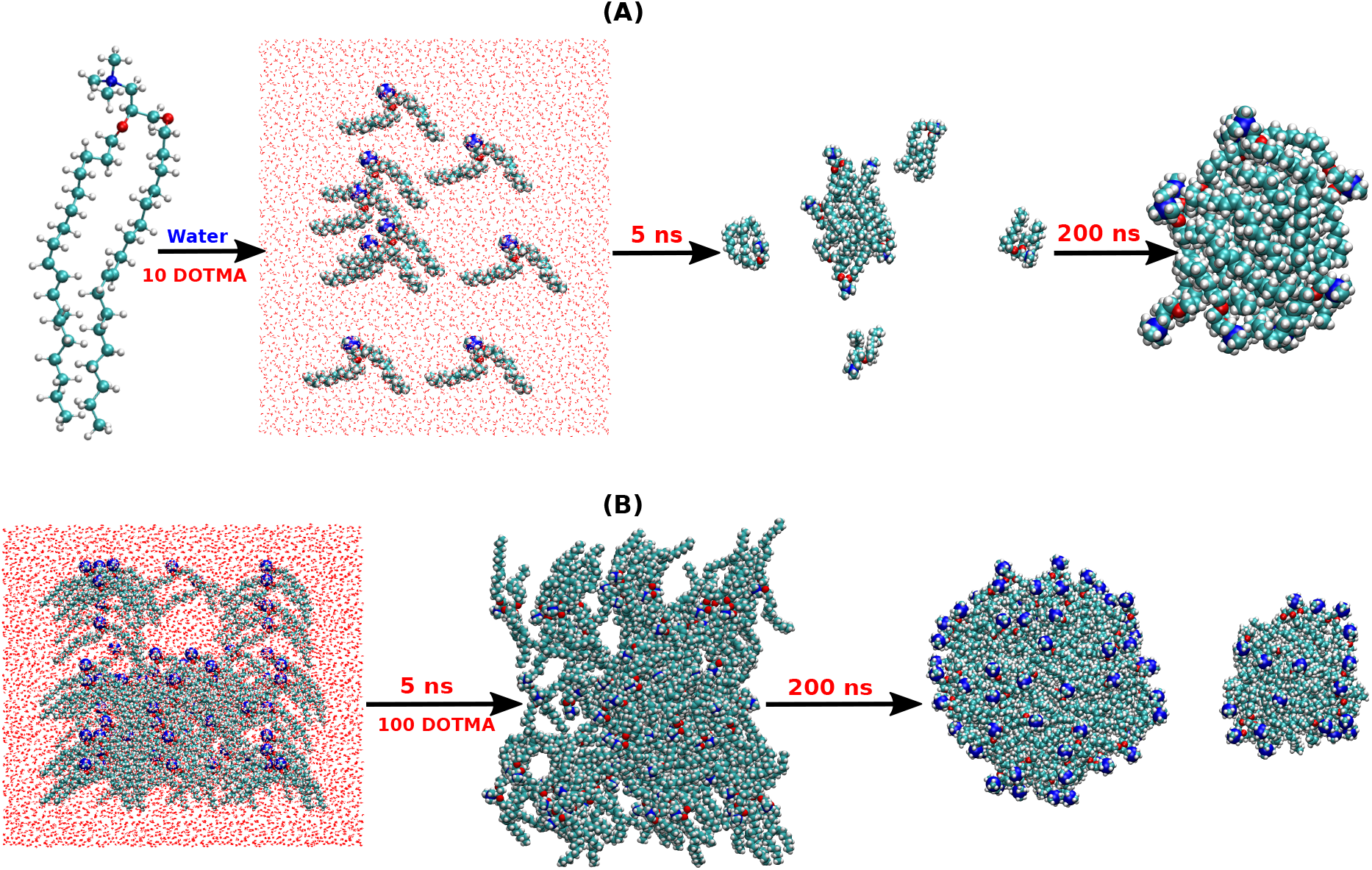
Representation of (A) cationic lipids DOTMA self-assembly: initial configuration of 10 DOTMA lipids (second), energy-minimized and equilibrated system in aqueous solution (third), and the final self-assembled cluster (fourth). (B) Initial and final morphologies of the cluster formation considering 100 cationic DOTMA lipids around the TDN nanocage, as shown by their instantaneous snapshots from the 200 ns MD simulation.

To order to investigate the distinct DOTMA clustering process with and without TDN, as shown in Figures 1 and 2, we have analyzed several structural properties. All the static structural properties were calculated from the last 50 ns of the 200 ns simulations. where the system reached conformational steady state and completed the TDN-DOTMA complexation process. The key morphological features of the formed clusters, such as average size (R_g_), mass, charge distribution, and the spatial arrangement, were characterized using radial distribution functions. The lipid clusters were identified by calculating the center of mass (COM) of each DOTMA lipid, and the lipids with COM distances *<*15 Å were grouped into the same cluster using the neighbor-based approach.

#### 1. Root-Mean-Square-Deviation (RMSD)

To assess the stability of the TDN cage as a potential drug delivery vehicle during interaction with DOTMA lipid clusters, we have calculated the root mean square deviation (RMSD) of the TDN cage with respect to an energy-minimized initial reference structure over 200 ns of MD simulations. As illustrated in Figure 3, the TDN nanocage exhibits stable RMSD profiles for different systems bound to 1, 10, or 100 DOTMA clusters, showing minimal deviation with respect to the bare TDN structure. This structural consistency underscores the stability of the TDN nanostructures and highlights the potential of DOTMA-functionalized TDNs as reliable candidates for drug delivery applications.

**FIG. 3.**
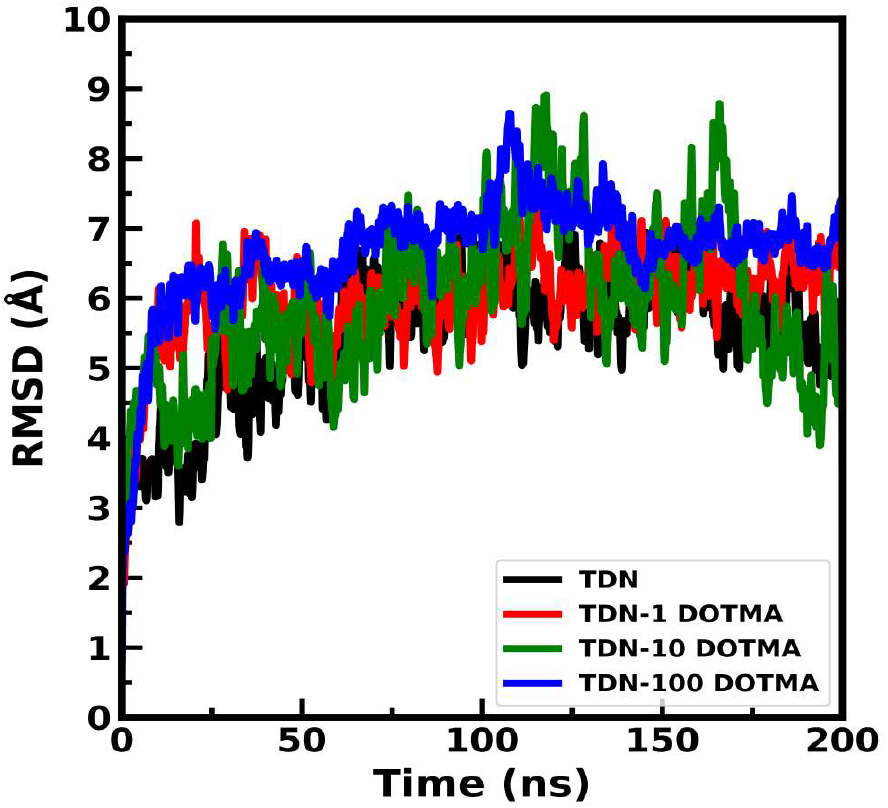
Time evolution of root-mean-square deviation of TDN cage in water and in the presence of 1, 10, and 100 DOTMA cationic lipids.

#### 2. Radial Distribution Function

To probe the spatial organization of DOTMA lipids and their interaction with TDN, we computed the radial distribution function (RDF) considering the stable part of the MD trajectory. The DOTMA lipids tend to self-assemble due to the hydrophobicity of the long lipid chains, while they are also expected to associate with the negatively charged TDN through strong electrostatic interactions with their cationic ammonium head groups. The tug-of-war between these two trends essentially drives their spatial equilibrium arrangements in the presence of TDN.

To elucidate the associative interactions, time-averaged RDFs were calculated between DOTMA-DOTMA and TDN-DOTMA complexes. For TDN-100 DOTMA complexation, RDFs between P-N1 atom pairs reveal a sharp peak near ~ 4.5 Å as shown in Figure 4A, indicating a strong electrostatic attraction and close binding propensity (also supported by minimum COM distance between TDN-DOTMA as discussed in section S.2 in the supplementary material). In contrast, the 1 and 10 DOTMA systems show low-intensity P-N1 RDF peaks, reflecting weaker and more dispersed TDN-DOTMA associative interactions. Figure 4B and C, shows that DOTMA lipids exhibit tighter packing in the presence of the TDN cage, as evidenced by sharp and high intensity RDF peaks near ~ 7.5 Å for O1–O1 and ~ 6 Å for N1–N1 atom pairs of DOTMA (as shown by black curves). In contrast, DOTMA–DOTMA shows comparatively weaker interactions without TDN, with less intense RDF peaks at similar distances. These observations suggest that the TDN promotes a more compact organization of DOTMA lipids, likely due to enhanced electrostatic interactions and the spatial confinement of lipids within the TDN cavity.

**FIG. 4.**
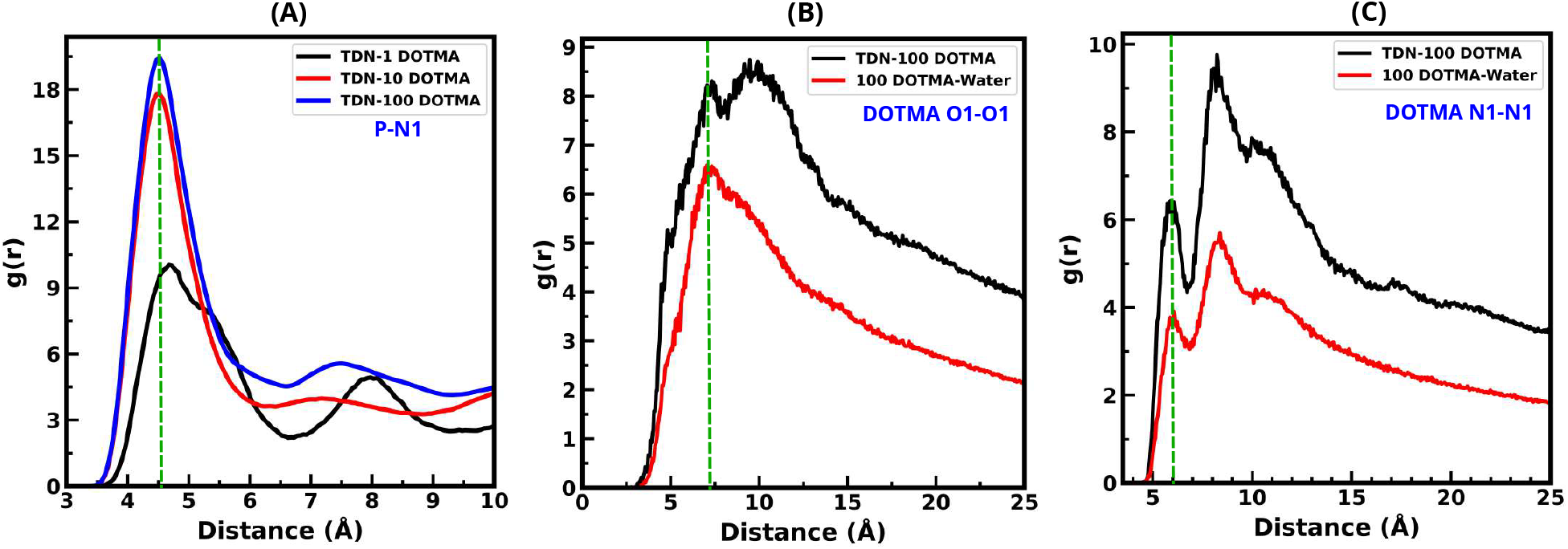
Radial distribution functions computed over the last 50 ns of the MD trajectory: (A) between TDN phosphate (P) atoms and DOTMA N1 headgroups for 1, 10, and 100 lipids, (B) between the O1-O1 atom pairs of DOTMA lipids both in presence and absence of TDN, (C) between N1-N1 atom pairs of DOTMA-DOTMA lipids with and without TDN, and. The description of O1 and N1 atoms is discussed in section S.1 in the supplementary material.

#### 3. Density Profiles

In order to characterize the distinct morphological features of the self-assembled DOTMA clusters both in the presence and absence of TDN, we investigated the mass and charge density profiles of only DOTMA clusters relative to their geometric center, as shown in Figure 5A–D. For 10 DOTMA lipids, both in the presence and absence of TDN, we observe a pronounced central mass density peak and a plateau extending up to 12 Å, indicating a compact core structure (Figure 5A). In contrast, with 100 DOTMA lipids in the presence of TDN (Figure 5B), the mass density peaks shift towards the cluster periphery with a central minimum. This shift suggests that the negatively charged TDN drives the heavier cationic N1 groups of DOTMA outward, forming a compact assembly with a dimension of ~ 70 Å (green curve). Conversely, in the absence of TDN, the 100 DOTMA lipids exhibit a central density peak with a gradual decay extending to 40 Å, which indicates a less compact DOTMA self-assembly in bulk water, spanning about 80 Å in cluster dimension (blue curve) as shown in Figure 5B. Figure 5C shows that for 10 DOTMA lipids, positive charge accumulates near the cluster surface (~ 15 Å) in both the presence and absence of TDN, followed by a sharp drop. This indicates a nearly spherical co-assembly with surface-localized positive charges, enabling rapid binding to the negatively charged TDN, though full encapsulation of the TDN cavity is limited by this small cluster of 10 lipids.

**FIG. 5.**
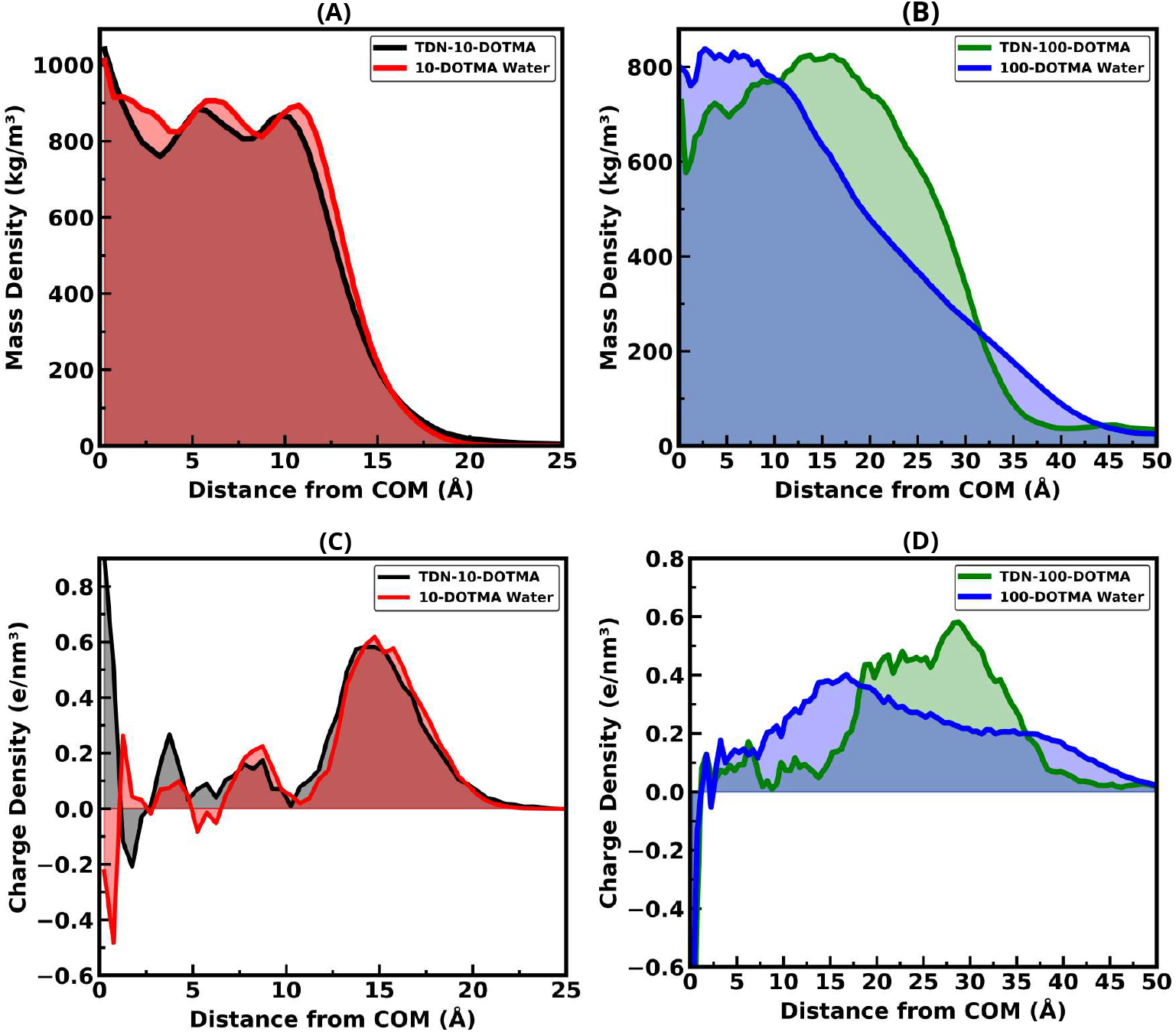
Mass density profiles for only DOTMA cluster: (A) 10 DOTMA with and without TDN cage, (B) 100 DOTMA with and without TDN cage. Charge density profiles for only DOTMA cluster: (C) 10 DOTMA in the presence and absence of TDN, (D) 100 DOTMA in the presence and absence of TDN, respectively.

For 100 DOTMA lipids without TDN, the partial charges are more uniformly distributed throughout the cluster with a low intensity peak around ~ 17 Å, followed by a uniform decline as shown in Figure 5D (blue curve). In contrast, in the presence of TDN, the charge distribution peaks near the cluster periphery (~ 30Å) with a central minimum, indicating that strong P–N1 electrostatic interactions drive the positively charged N1 head groups outward toward the negatively charged TDN surface, resulting in a complete entrapment of the compact DOTMA cluster within the TDN cavity (green curve).

#### 4. Radius of Gyration

DOTMA lipids self-assemble into a stable cluster starting from a randomly dispersed state, both in the presence and absence of the TDN cage. We estimated the size of the self-assembled lipid cluster, as well as the spatial dimensions of the individual DOTMA lipids and TDN cage, using the average radius of gyration.

The radius of gyration (R_g_) of individual DOTMA lipids remains nearly unchanged across different DOTMA ratios (1, 10, and 100), both in the presence and absence of TDN cages, indicating minimal impact on lipid flexibility (compaction or expansion of the shape or conformation) as shown in Figure 6A. The morphological evolution of self-assembled DOTMA clusters was also analyzed by tracking their average R_g_. The time evolution of average cluster R_g_ shows that for the 10 DOTMA system, the cluster size in bulk water is near 12.97±0.32 Å, while in the presence of TDN, it slightly decreases to 11.59±0.17 Å, suggesting more compact packing due to electrostatic interactions as shown in Figure 6B. This implies similar, nearly spherical final morphologies regardless of TDN.

**FIG. 6.**
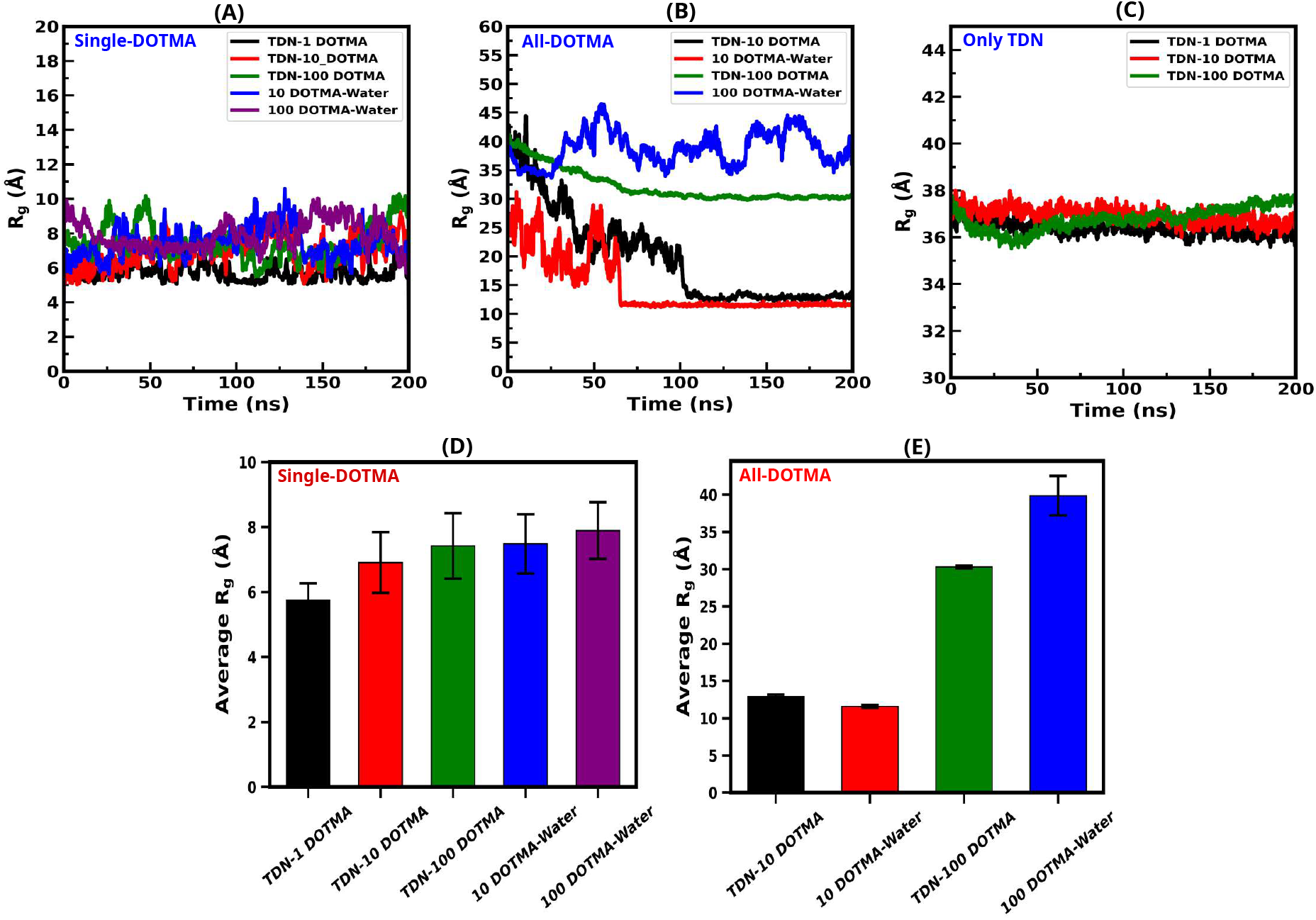
Radius of gyration plots are shown for: (A) individual DOTMA lipids across all five systems, (B) considering all cluster-forming DOTMA lipids, (C) only TDN nanocage, (D–E) time-averaged R_g_ values over the last 50 ns.

However, the 100 DOTMA system exhibits notable differences in cluster size in both with and without TDN. A comparable trend is observed for the 100 DOTMA system, as shown in Figure 6B. In the presence of TDN, strong electrostatic interactions lead to greater lipid compaction, resulting in lower R_g_ values (green curve). The lipid cluster fully occupies the TDN cavity, forming well-defined, nearly spherical co-assemblies. In contrast, clusters in the bulk water show higher fluctuating R_g_ of *~*37 Å due to looser packing, which decreases to ~ 34 Å in the presence of TDN cage, due to tighter entrapment in the cavity. Additionally, the R_g_ of TDN cages remains stable across different lipid ratios, indicating structural integrity of TDN nanocage upon DOTMA binding as shown in Figure 6C–E, also supported by the RMSD results.

#### 5. Effect of Hydration in the DOTMA Binding

Morphological and structural analyses show that 100 DOTMA lipids first self-assemble and the corresponding lipid cluster becomes entrapped within the TDN cavity, where it interacts and associates with the negatively charged TDN backbones, neutralizing negative charges and potentially enhancing cellular uptake. To uncover the driving force behind this entrapment, we examined hydration of TDN nucleotides with different DOTMA ratios, focusing on the first and second hydration shells as shown in Figure S.3 in the supplementary material, with cutoffs distances of 3.5 Å for the first hydration shell and 5.0 Å for the second hydration shell. As shown in Figure S.3, a significant disruption in the hydration network occurs at the 100 DOTMA ratio, with reduced water density near TDN nucleotides, particularly inside the cavity. We hypothesize that this dehydration, combined with TDN’s pyramid-like topology, creates a hydrophobic environment that promotes DOTMA clusters’ encapsulation inside the TDN cavity.

To probe hydration dynamics of TDN cavity at varying lipid ratios, we have also investigated 2D density distribution maps of water and Mg^2+^ ions. As shown in Figure 7A–B, the TDN cavity remains well hydrated with 1 DOTMA lipid, but becomes dehydrated and fully occupied by the more hydrophobic DOTMA clusters composed of 100 lipids (Figure 7C–D). A corresponding drop in Mg^2+^ ion density within the TDN cavity is also observed as shown in Figure S.4 in the supplementary material, highlighting reduced hydration within the TDN cage relative to the exterior. These findings suggest that hydration plays a key role in driving the entrapment of DOTMA clusters inside the TDN cavity.

**FIG. 7.**
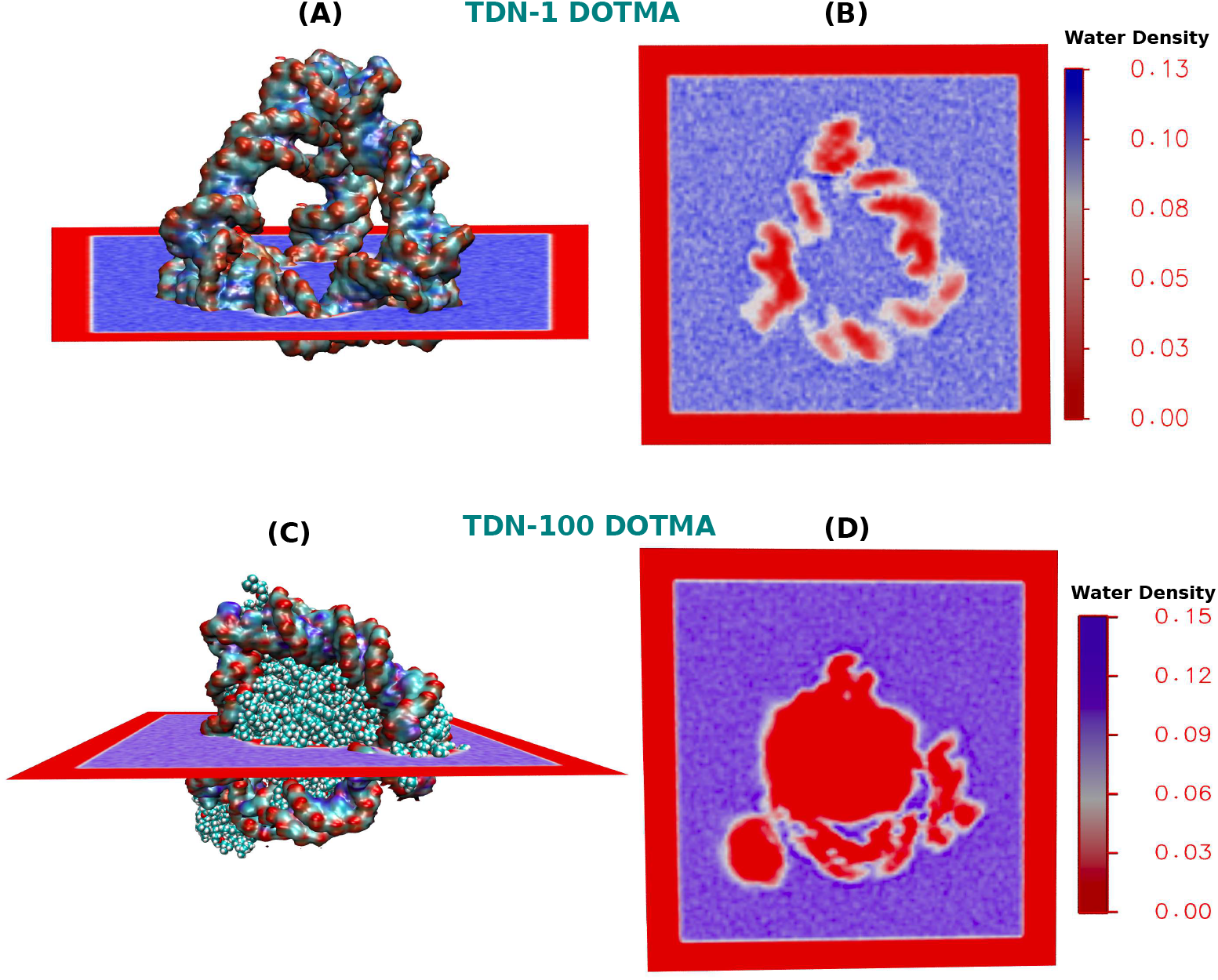
Representations of the volume density distribution maps of water molecules to get an overview of their hydration in the presence of DOTMA cationic lipids around the TDN nucleotides. The color bar represents the density distribution of water molecules in the units of atoms/Å^3^

### B. Energetic Properties

In the previous section, we discussed the morphology of self-assembled DOTMA-DOTMA and TDN-DOTMA structures. Here, we focus on how the interaction energies drive their association. A quantitative understanding of TDN–DOTMA complexation is obtained by estimating their non-bonded interaction energies and binding enthalpies, since there are no covalent bonds exist between them.

#### 1. Non-bonded Interaction Energies

To better understand the energetic contributions driving TDN-DOTMA complexation and DOTMA-DOTMA self-assembly in bulk water, we analyzed the relevant energy components. As there are no covalent bonds between DOTMA and TDN, their interactions are entirely governed by non-bonded forces, with electrostatic interactions consistently dominating over VDW contributions across all systems, as illustrated in Figure 8 and the quantitative values are listed in Table 1 in the SI.

**TABLE I.** Details of the five systems simulated in this work.

| System | # of TDN bases | # of DOTMA Lipids | Conc. of $\text{MgCl}_2$ (mM) | # of Neutralizing Atoms ($\text{Mg}^{2+}$) | # of Water Molecules | # of Total Atoms |
| --- | --- | --- | --- | --- | --- | --- |
| TDN + 1 DOTMA | 240 | 1 | 150 | 120 | 59529 | 186718 |
| TDN + 10 DOTMA | 240 | 10 | 150 | 115 | 72680 | 227381 |
| TDN + 100 DOTMA | 240 | 100 | 150 | 70 | 74269 | 242891 |
| Only 10 DOTMA | - | 10 | 150 | - | 16082 | 49612 |
| Only 100 DOTMA | - | 100 | 150 | - | 41490 | 137544 |

**FIG. 8.**
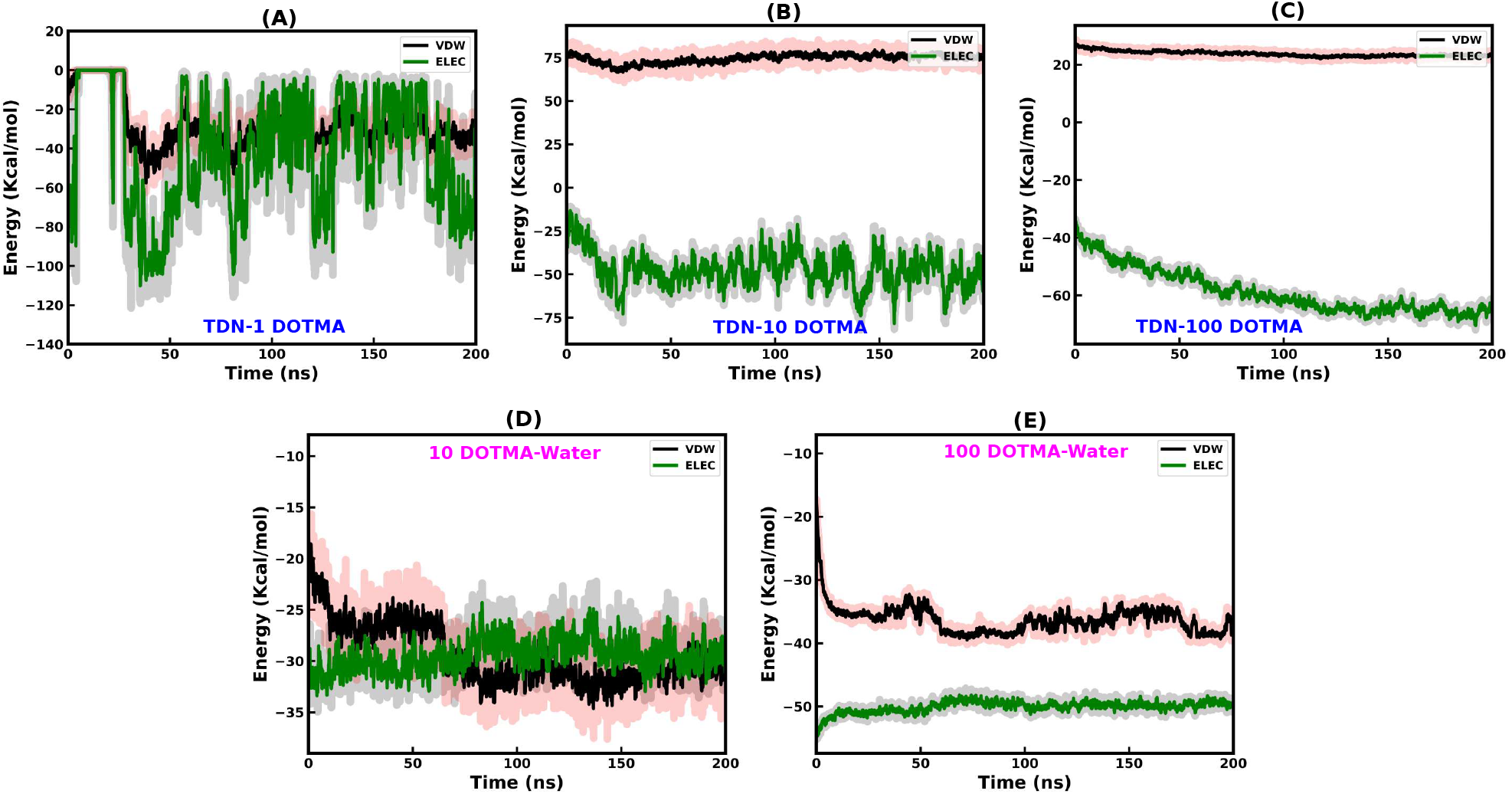
Time evolution of the non-bonded interaction energies per-DOTMA; van der Waals (VDW, black) and electrostatic (green) for the following systems: (A) TDN–1 DOTMA complex, (B) TDN–10 DOTMA complex, (C) TDN–100 DOTMA complex, (D) 10 DOTMA-DOTMA lipids in bulk water, and (E) 100 DOTMA-DOTMA lipids in bulk water

Figure 8A-C shows the time evolution of non-bonded energy per-DOTMA lipid for all three TDN–DOTMA systems. In the TDN-1 DOTMA system, electrostatic energy initially fluctuates due to the absence of the DOTMA clusters and binds to the outer mode of TDN with a binding strength of ~ - 24 kcal/mol (detailed energy component values are listed in Table 2, SI). Similarly, in the TDN-10 DOTMA system, electrostatic energy per-DOTMA dominates over VDW energy, with stable binding of the DOTMA cluster achieved near ~ 50 ns. For the 100 TDN-DOTMA system, electrostatic energy increases significantly during the first 150 ns, corresponding to the hydrogen bond formation between TDN-DOTMA and their complex stabilization. Beyond 150 ns, the electrostatic energy plateaus (green curve), indicating the most stable binding state with 100 DOTMA lipids, among the three systems (Figure 8C) in the presence of TDN. The quantitative estimates of the MMGBSA binding energy values for the three TDN-DOTMA complexes are listed in Table 3 in the SI.

In contrast, DOTMA–DOTMA self-assembly in bulk water, with 10 and 100 DOTMA systems shows a rapid electrostatic stabilization within ~ 10 ns, driven by rapid lipid clustering facilitated by both hydrophobic and electrostatic interactions in the absence of TDN (Figure 8D-E). However, the electrostatic energy is more stable but lower in magnitude for DOTMA-DOTMA self-assembly compared to TDN-DOTMA complexation, which highlights the fact that in the absence of TDN, DOTMA lipids quickly self-assemble, whereas in the presence of TDN, the tug-of-war between lipid self-assembly and TDN-DOTMA complexation delays electrostatic stabilization as shown in Figure 8C.

#### 2. Electrostatic Potential Maps

To get insights into the electrostatic interactions driving the complexation of TDN-DOTMA conjugates, we computed the electrostatic potential distribution map *φ* (**r**) for the TDN-DOTMA complex by solving the Poisson equation:

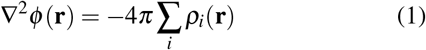

where the summation runs over all charged atoms of TDN-DOTMA complex, and the charge density *ρ*_*i*_(**r**) for each atom is approximated using a spherical Gaussian distribution:

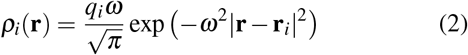

Here, *q*_*i*_ is the partial charge of the *i*-th atom, **r**_*i*_ is its position, and *ω* is the inverse width of the Gaussian distribution^44^. The instantaneous electrostatic potential *φ* (**r**) was averaged over the final 20 ns of the simulation trajectory, accounting for all charged atoms in the system.

The electrostatic potential distribution map reveals how the negative charges of TDN backbone are effectively neutralized through non-covalent complexation with cationic DOTMA lipids, thereby enhancing the conjugate’s ability to permeate negatively charged cellular membranes. In the TDN–1 DOTMA system, the TDN structure retains a pronounced negative electrostatic potential distribution, as indicated by deep violet regions in Figures 9A–B, which likely hinders membrane permeability due to electrostatic repulsion. In contrast, Figures 9C–D illustrate that the presence of 100 DOTMA lipids results in the formation of a compact lipid cluster that interacts extensively with the negatively charged TDN backbones through stronger electrostatics. This interaction significantly reduces the negative electrostatic potential of the complex, as observed through the prominent charge-neutral lighter violet regions, thereby potentially facilitating improved membrane permeation.

**FIG. 9.**
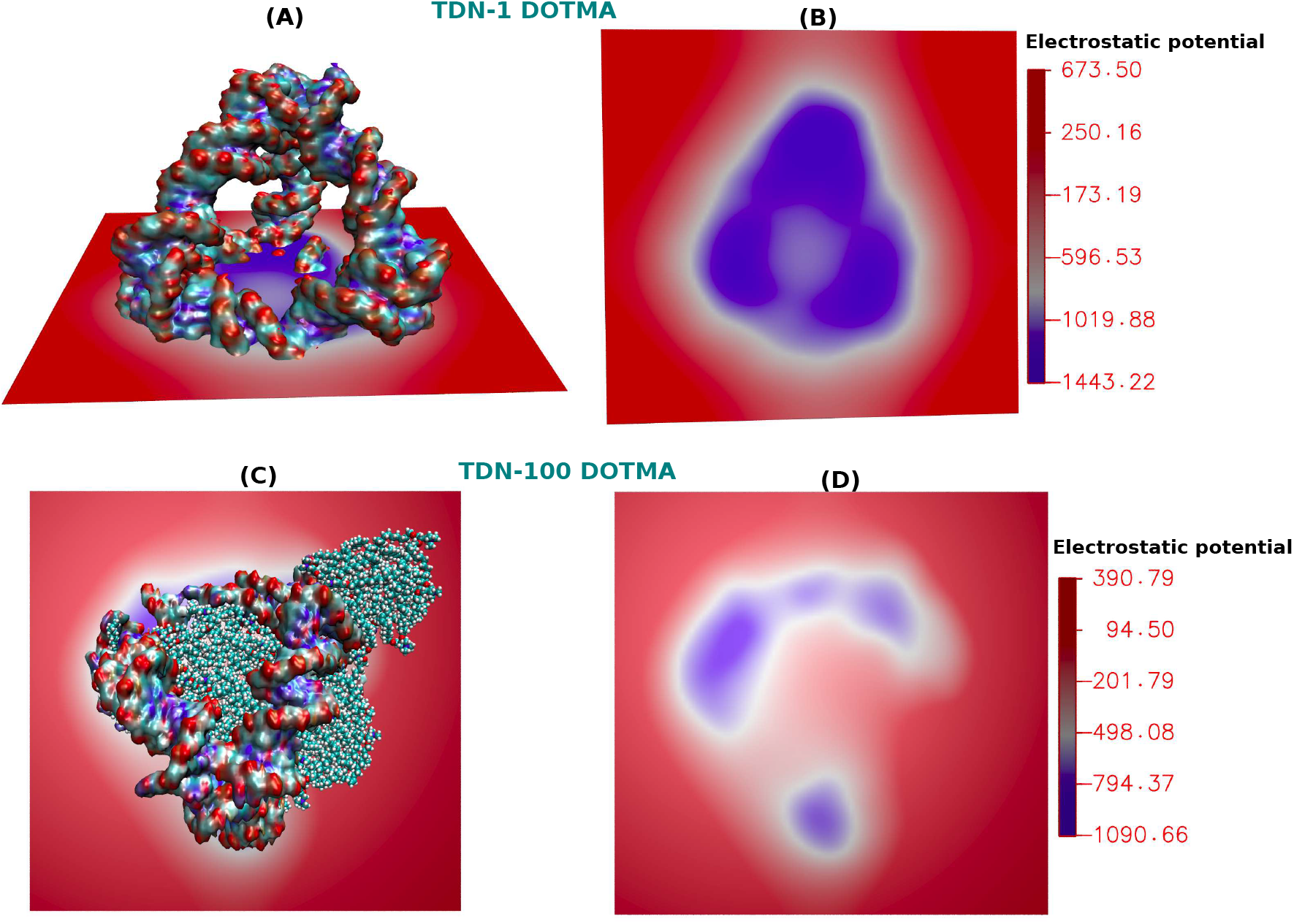
Electrostatic potential maps for the TDN-DOTMA systems containing 1 and 100 DOTMA lipids illustrate the extent of DOTMA binding, as seen by the reduced presence of deep violet regions across the planes. The X–Y plane provides a top-down view, while the Z–Y planes show the side perspectives of the potential distribution. In the TDN–100 DOTMA system, stronger binding of cationic lipids to the DNA backbone results in more effective neutralization of the negative potential, reflected by the expansion of white regions, indicating enhanced electrostatic screening. The color scale denotes the electrostatic potential in units of K_B_T/e

### C. Dynamic Properties

#### 1. Association Kinetics with Number of Clusters

We now focus on the time scale associated with the self-assembly kinetics of the cationic lipids, a key factor in designing lipid-based nanoparticles for efficient nucleic acid delivery and transfection. To explore this, we analyzed the time evolution of lipid cluster formation, starting from an out-of-equilibrium state where DOTMA lipids were randomly distributed in water and also around the TDN, to an energetically stable state. As shown in Figure 10A, DOTMA clustering kinetics vary with lipid ratios and also in the presence of TDN. Interestingly, Figure 10A shows that in presence of TDN, most of the DOTMA lipids form a compact cluster inside the TDN cavity, but the process is relatively slower as compared to their cluster formation in bulk water due to competing effects of hydrophobicity, hydrogen bonding, and electrostatic interactions. Absence of this tug-of-war seems to result in a rapid clustering for the 10 and 100 DOTMA lipids in bulk water within the first 50 ns.

**FIG. 10.**
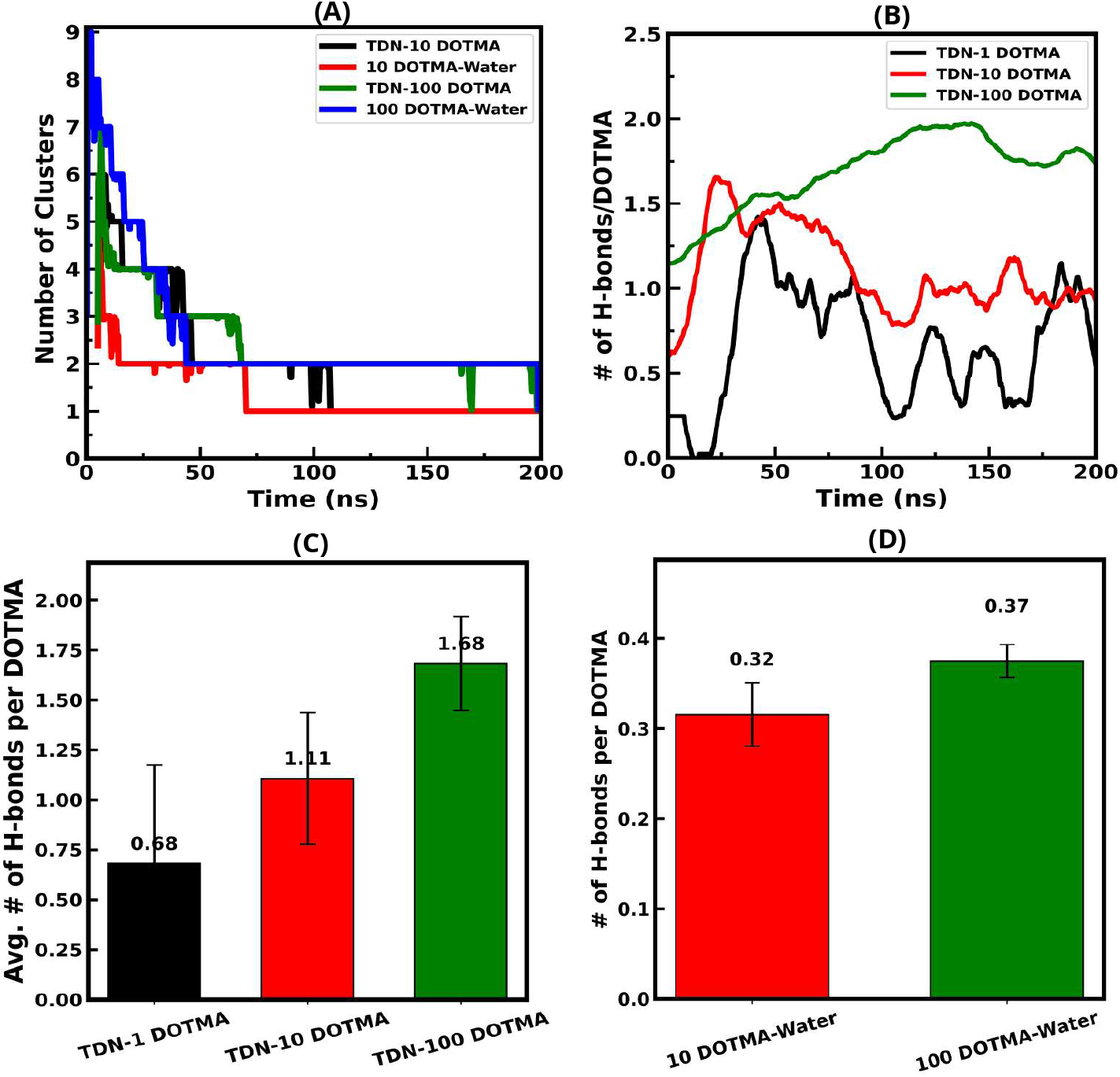
(A) Time evolution of the number of DOTMA cluster distributions formed across different lipid ratios. (B) Number of hydrogen bonds per-DOTMA lipid formed between DOTMA and TDN over the simulation time. (C–D) Average number of hydrogen bonds per-jDOTMA lipid between TDN–DOTMA and DOTMA–DOTMA interactions, calculated over the final 50 ns of the simulation, both with and without the presence of TD

#### 2. Hydrogen Bonding Dynamics

Hydrogen bonding plays a key role in the association kinetics of TDN and DOTMA lipids, as an increasing function of time, as the approach of the two components evolves over time. Using a distance cutoff of 3.5 Å and an angle cutoff of 50 degrees, we quantified number of hydrogen bonds per-DOTMA lipid over time as shown in Figure 10B.

The number of hydrogen bonds per-lipid for the TDN-100 DOTMA system initially increases up to 150 ns and then attains a steady state. Further restructuring of the lipids within the clusters after complexation with TDN justifies the fluctuation of the curves at the long-time limit after 150 ns. However, the large drop in the number of cluster distribution curves in Figure 10A (green curve), at short timescales (within 100 ns) indicates that the cluster formation precedes their association with TDN, since hydrogen bonding between TDN and lipid molecules still continues to increase at these times up to ~ 150 ns as shown in Figure 10B. For 1 and 10 DOTMA lipids, per-DOTMA hydrogen bonding with TDN fluctuates due to weaker binding affinity with one of the TDN edges and the dynamic nature of hydrogen bonding during the simulation.

The time-averaged hydrogen bonds for 1, 10, and 100 DOTMA lipids binding to TDN nucleotides, as well as between 10 and 100 DOTMA-DOTMA lipids in bulk water, are shown in Figure 10C-D, clearly indicating that the 100 DOTMA lipid ratio is crucial for stronger association with the considered TDN nanocage. As shown by the green bar in Figure 10C, the number of hydrogen bonds per-DOTMA becomes maximum due to stronger electrostatic attraction in the presence of TDN and 100 DOTMA lipids. In contrast, the only DOTMA–DOTMA hydrogen bonds remain relatively constant as shown in Figure 10D, reflecting stable clustering of hydrophobic lipids in water.

#### 3. Dynamic Radial Distribution Function

To investigate the temporal evolution of the TDN-DOTMA complexation and lipid aggregation process, we analyzed radial distribution functions at different time intervals. Figure 11A-C presents dynamic RDFs, to capture the dynamic interactions between TDN phosphate (P) and DOTMA N1 atoms, as well as between N1-N1 atom pairs of the DOTMA lipids.

**FIG. 11.**
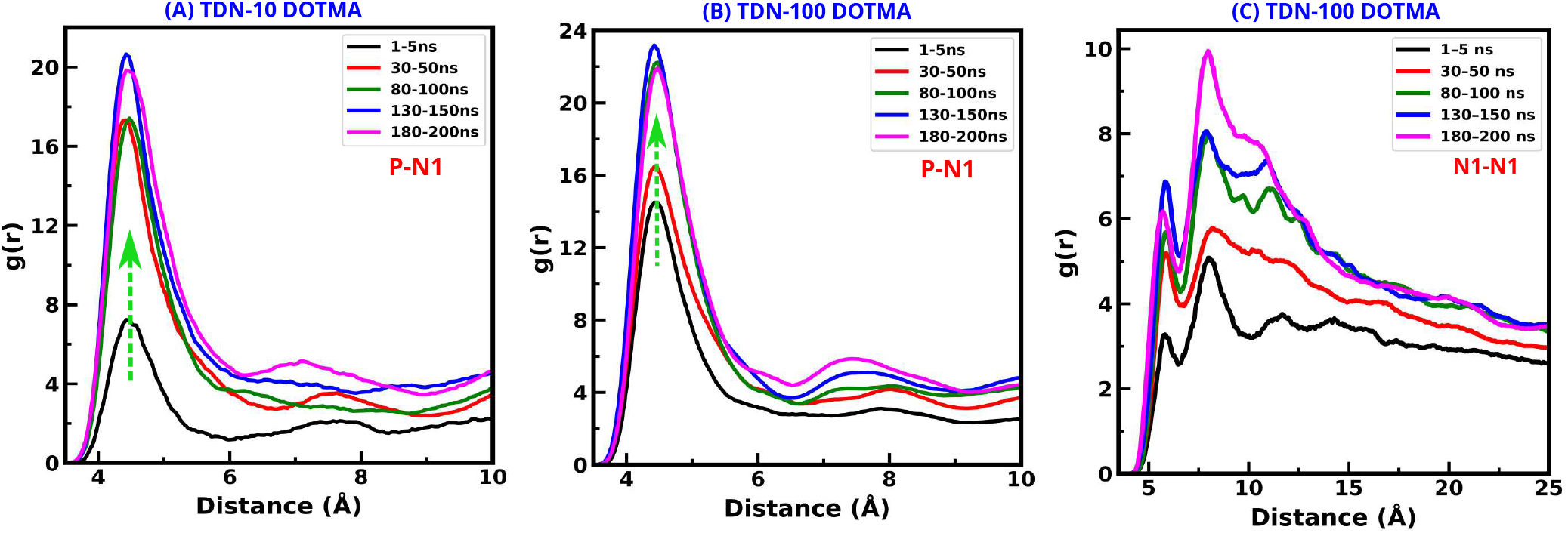
Radial distribution function (RDF)- (A, B) between the phosphate (P) atoms of TDN nucleotides and N1 atoms of the 10 and 100 DOTMA lipids at different time intervals to evaluate electrostatic association dynamics, (C) between N1-N1 atoms pairs of DOTMA lipids for 100 DOTMA systems. The 100 DOTMA ratio is the most significant one, as it shows complete entrapment in the TDN cavity and nearly spherical co-assembly as observed in simulation as well as in experiment.

In the TDN-10 DOTMA system, P-N1 RDF shows a rapid increase of the first peak between 30-50 ns, which indicates TDN-DOTMA complexation as the self-assembled DOTMA clusters bind to the TDN, as shown in Figure 11A. Thereafter, a gradual enhancement of the P-N1 RDF peak in terms of intensity and width takes place, until a convergence is observed beyond 50 ns, highlighting the formation of a stable TDN-10 DOTMA conformational state.

In the TDN–100 DOTMA system, sharp rise of the P-N1 RDF peak near 80-100 ns, from the randomly distributed initial state indicates complexation of lipids with TDN, but at later times sharper and converging P–N1 RDF peak beyond 100 ns, indicating the formation of a stable TDN–DOTMA complex as shown in Figure 11B. Therefore, in the presence of TDN, 100 DOTMA lipids first self-assemble into a DOTMA cluster within 70 ns (Figure 10A), and only then associate with TDN, confirming that self-assembly precedes the TDN-DOTMA complexation.

Figure 11C presents the timescale of DOTMA-DOTMA self-assembly within the TDN–100 DOTMA system by tracking the dynamic RDF of N1–N1 atom pairs. A sharp increase of N1-N1 RDF peak near 30 ns (red curve) indicates rapid self-assembly of the lipids within the TDN cavity. Beyond 30-50 ns, this peak quickly converges to higher intensity, consistent with the self-assembled cluster distribution timescale (*~*70 ns). These results further support that DOTMA self-assembly occurs prior to TDN-DOTMA complexation, as evidenced by the RDF convergence times

We simulated the TDN cage with 250 DOTMA lipids at physiological salt concentration and observed initial cluster formation followed by complexation with the TDN. Unlike the nearly spherical co-assembly observed with 100 DOTMA lipids, the final morphology with 250 DOTMA forms a network-like structure (Figure S.5, SI), as also observed in the experiment^20^.

## IV. CONCLUSION

Using extensive atomistic MD simulations, we have investigated the molecular origins of the complexation mechanism of cationic DOTMA lipids with negatively charged phosphate groups of TDN in a physiological environment. This has been motivated by the experimental evidence of enhanced cellular uptake with DOTMA-conjugated DNA nanostructures through cellular plasma membranes^20^. Our results show that DOTMA lipids form structurally and energetically stable clusters both with and without TDN. Charge distribution profiles reveal that in 100-DOTMA cluster in the presence of TDN, positive charges localize at the periphery, maximizing electrostatic interactions. Radius of gyration analysis indicates a compact, stable 100 DOTMA lipids co-assembly in presence of TDN, whereas DOTMA-DOTMA clusters are comparatively less compact. Overall, DOTMA lipids rapidly self-assemble within the first few nanoseconds, and we capture key factors driving both DOTMA self-assembly and TDN–DOTMA complexation.

We observed two distinct phenomena with increasing DOTMA concentration in the aqueous solution: (a) First, rapid formation of self-assembled clusters of DOTMA lipids driven by strong hydrophobic interactions, (b) Secondly, slower TDN-DOTMA complexation, where clusters become entrapped within the TDN cavity, primarily driven by electrostatics, hydrogen bonding, and hydration of the TDN. These findings confirm that DOTMA self-assembly precedes complexation with TDN. We observe that at a high lipid ratio (100 DOTMA), the non-covalent functionalization results in complete entrapment of the DOTMA cluster within the TDN cavity, which leads to strong P–N1 interactions and effective neutralization of TDN’s negative charge, thereby enhancing its drug delivery potential.

This study provides new mechanistic insights into molecular understanding of the non-covalent conjugation of cationic lipids with DNA nanostructures to form a stable complex, which will shed light on the future development of lipid-functionalized DNA nanostructures for optimized drug delivery/transfection vehicles, bioimaging, and various biomedical applications. In the case of gene delivery, this variant of the TDN-DOTMA complex may show promise in offering defense, for example, against enzymatic degradation. This work also laid the foundation for developing RNA- and other nucleotide-based nanocarriers functionalized with cationic lipids for enhanced cellular internalization.

## SUPPLEMENTARY MATERIAL

See the supplementary material for a detailed description of structural and thermodynamic properties, which supports the information provided in the main manuscript.

## ACKNOWLEDGMENTS

SM acknowledges the SRF fellowship from CSIR, India. PKM thanks the Department of Science & Technology (DST), India, for financial support and the Science and Engineering Research Board (SERB) for financial and computational support through CRG/2021/003659.

## AUTHOR DECLARATIONS

### Ethics Approval

Ethics approval is not required.

## DATA AVAILABILITY

The data that support the findings of this study are available from the corresponding author upon reasonable request

