## Supplementary Information (SI) for "In silico self-assembly and complexation dynamics of cationic lipids with DNA nanocages to enhance lipofection"

### S.1 DOTMA Conformation

The cationic lipid N-[1-(2,3-dioleyloxy)propyl]-N,N,N-trimethylammonium (DOTMA) was selected as a non-viral complexation agent through non-covalent functionalization with the nucleotides of TDN for better transfection efficiency/cellular uptake. Due to their pyramid-like structure and duplex DNA edges, TDN exhibits exceptional stiffness and stability, making it ideal for drug delivery. DOTMA consists of three important parts: (1) a cationic ammonium head that stabilizes DNA TDN via electrostatic attraction; (2) a glycerol backbone with ether linkages that improve transfection and biological half-life; and (3) hydrophobic hydrocarbon chains that facilitate plasma membrane permeation.

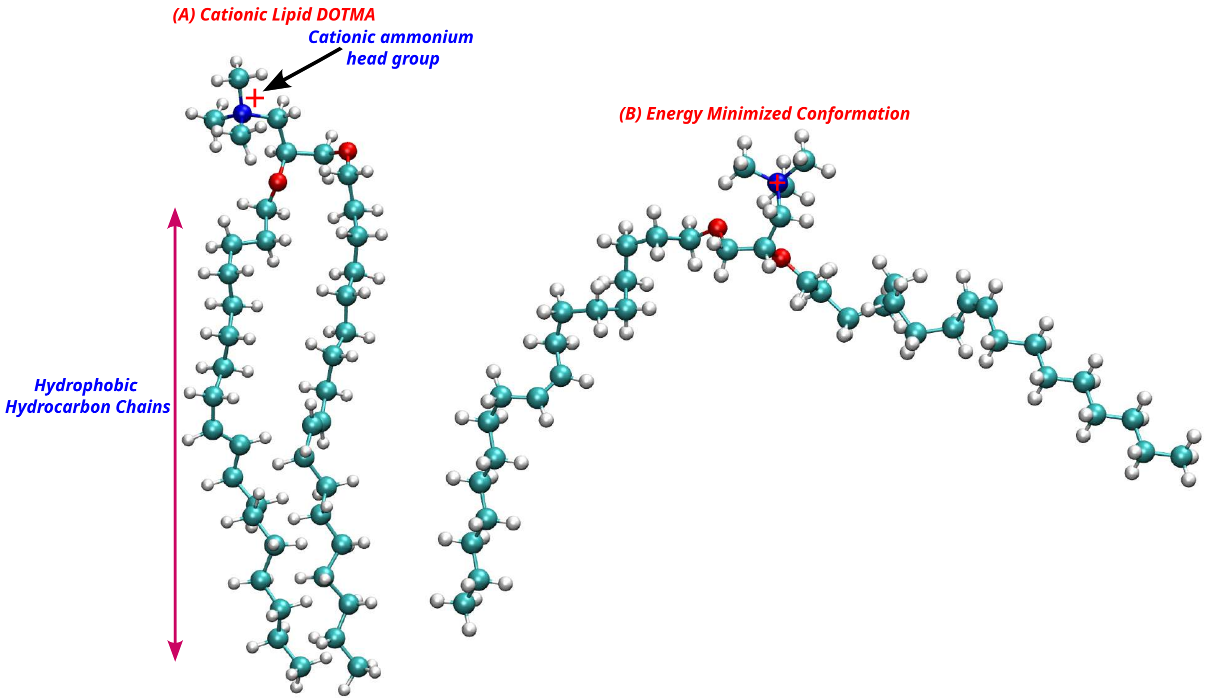


Figure S.1: Instantaneous snapshots of the initial and energy minimized DOTMA structures.

### S.2 Minimum Distances between TDN and DOTMA Lipids

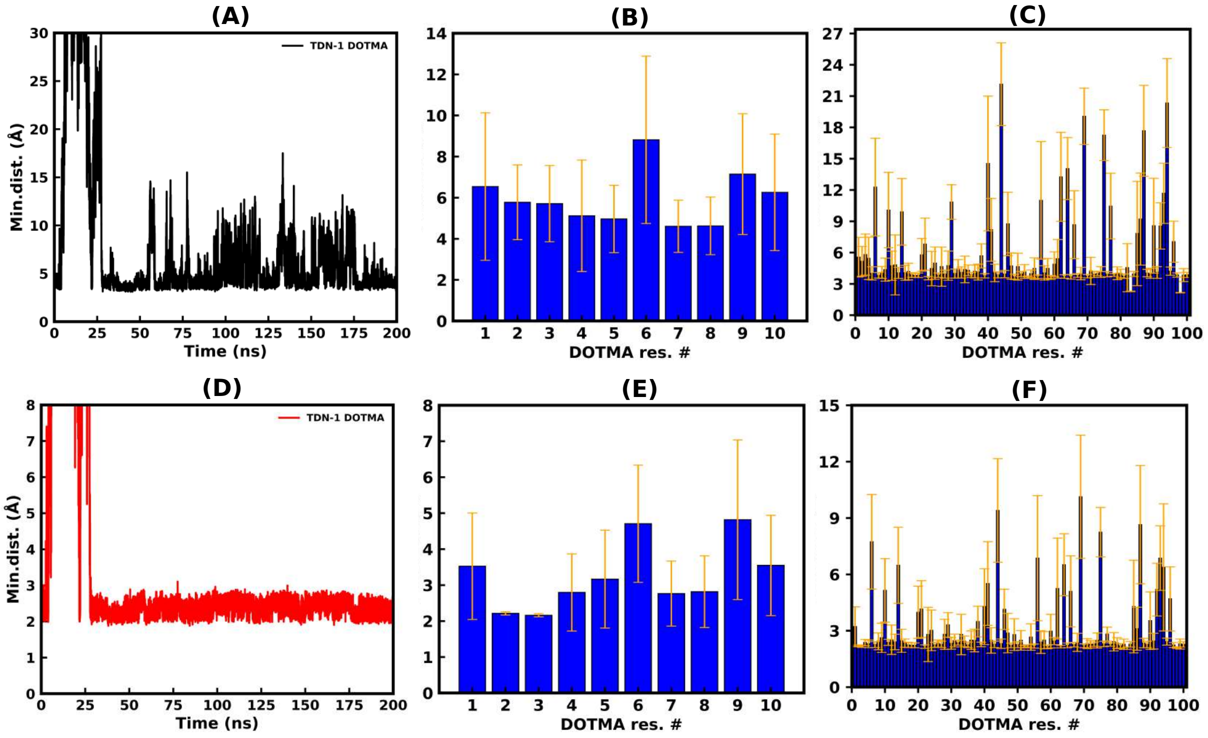


Figure S.2: Minimum distances between TDN nucleotides and N1 head groups of DOTMA for [A]1 DOTMA, (B) 10 DOTMA, and (C) 100 DOTMA, respectively. Also, the minimum distances between the center of mass of TDN nucleotide and DOTMA lipids for the 1, 10, and 100 lipids are represented by (D), (E), and (F), respectively.

### S.3 TDN Hydration shells

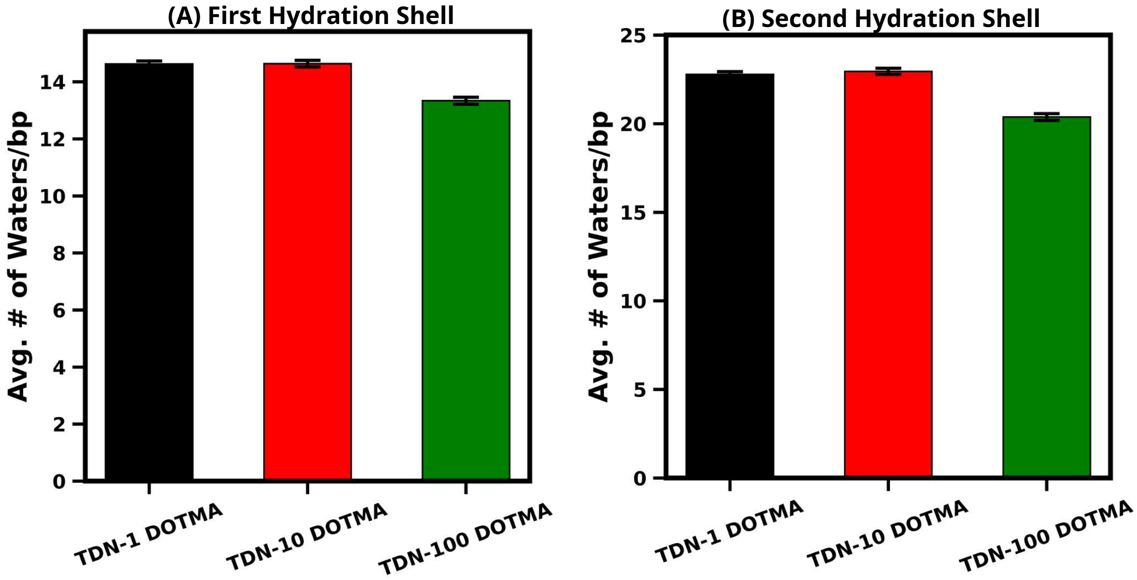


Figure S.3: *Representations of number of water molecules per base-pair for the first and second hydration shells near TDN nucleotides. (A) Number of water molecules per base-pair in first hydration shell within 3.5 Å, (B) Number of waters in second hydration shell with 5 Å*

### S.4 Hydration of MG ions around TDN cage

TDN cavity remains hydrated by waters and MG ions for the case of TDN-1 DOTMA lipid but becomes dehydrated and filled with hydrophobic DOTMA clusters for 100 lipids. This is accompanied by a marked decrease in $Mg^{2+}$ density inside the TDN, indicating reduced internal hydration.

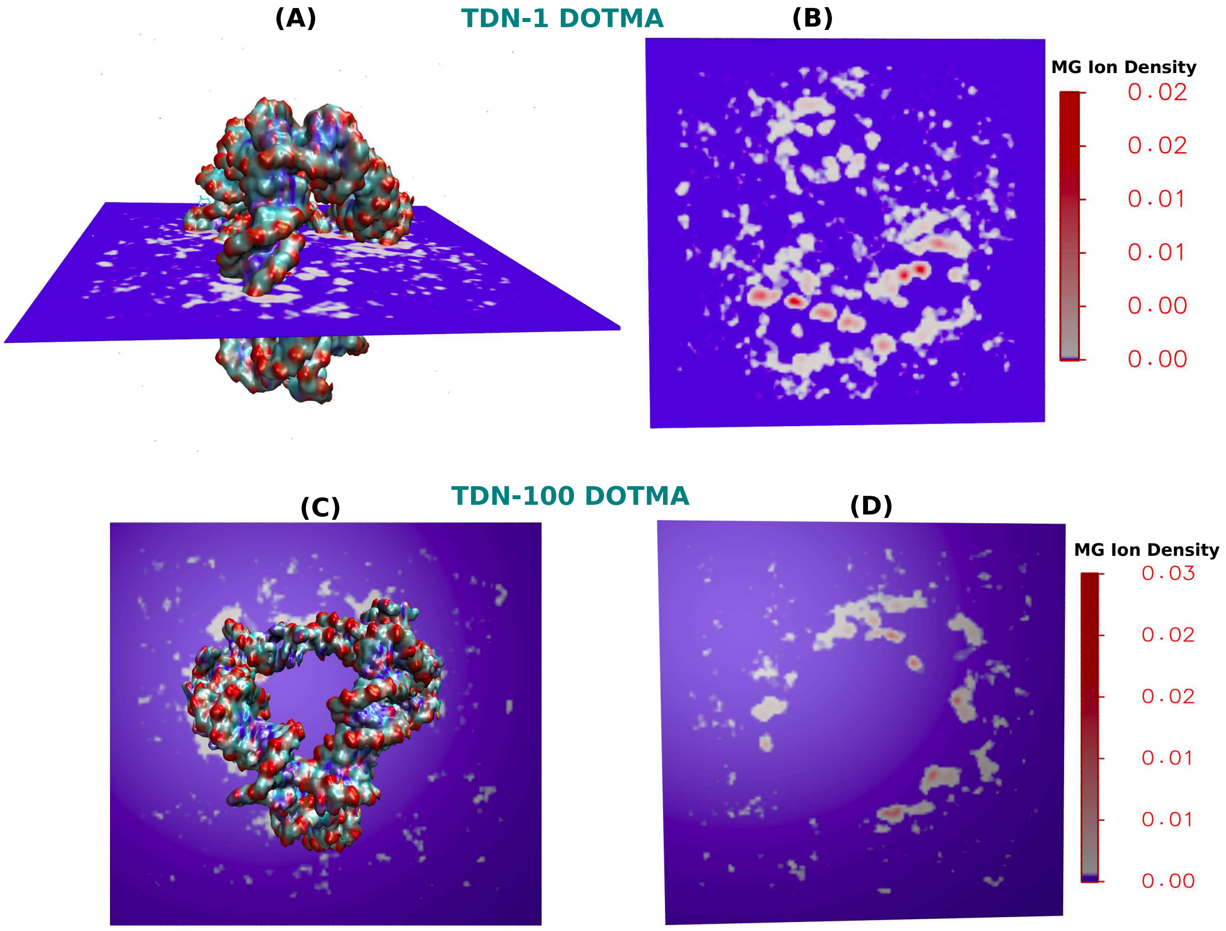


Figure S.4: *Representation of volume density maps- (A-B)top and side view for MG ions density for the TDN with 1 and 100 DOTMA lipid systems at 150 mM salt concentrations, respectively. The color bar represents the MG ions density in atoms/Å*$^3$

### S.5 Time-averaged total non-bonded interaction energies

Table St.1: Average total non-bonded interaction energies between DOTMA and TDN nucleobases, separated into van der Waals and electrostatic components, calculated over the final 50 ns of the 200 ns simulation.

| **System** | $<\mathbf{E_{ELE}}>$ | $<\mathbf{E_{VDW}}>$ | $<\mathbf{E_{Total}}>$ |
|---|---|---|---|
| TDN+1 DOTMA | -41.53±26.83 | -31.26±4.76 | -72.80±29.57 |
| TDN+10 DOTMA | -497.51±89.42 | 757.65±23.95 | 260.14±100.58 |
| TDN+100 DOTMA | -6515.97±128.67 | 2320.35±53.21 | -4195.62±202.76 |
| 10 DOTMA without TDN | -294.81±15.35 | -307.89±15.40 | -602.71±18.96 |
| 100 DOTMA without TDN | -4961.44±56.31 | -3766.43±128.42 | -8727.87±117.70 |

### S.6 Binding Free Energy using MMGBSA Approach

The binding free energy calculation was performed to determine the interaction strength of association between the TDN nucleotides and cationic DOTMA lipids. The MMGBSA method in the MMPBSA.py[1–5] module of AMBER20[6] was used to calculate the binding free energy of TDN-DOTMA complex formation as follows:

$$\Delta G_{\text{binding}} = \Delta E_{\text{complex}} - \Delta E_{\text{TDN}} - \Delta E_{\text{DOTMA}} \tag{1}$$

Where $\Delta E_{\text{complex}}$, $\Delta E_{\text{TDN}}$ and $\Delta E_{\text{DOTMA}}$ represent free energies of the TDN-DOTMA complex, individual TDN, and individual DOTMA receptors, respectively. The eqn-1 can be decomposed into different interactions and can be written as -

$$\Delta G_{\text{binding}} = \Delta H - T\Delta S \simeq \Delta E_{\text{MM}} + \Delta G_{\text{sol}} - T\Delta S \tag{2}$$

Here, $\Delta E_{\mathrm{MM}}$ is the change in gas-phase molecular mechanics energy, $\Delta S$ is the entropy change, and $\Delta G_{\mathrm{sol}}$ represents the solvation free energy change upon ligand binding. $\Delta E_{\mathrm{MM}}$ can be further written as-

$$\Delta E_{\mathrm{MM}} = \Delta E_{\mathrm{bonded}} + \Delta E_{\mathrm{vdW}} + \Delta E_{\mathrm{elec}} \tag{3}$$

Here $\Delta E_{\mathrm{bonded}}$, $\Delta E_{\mathrm{elec}}$, and $\Delta E_{\mathrm{vdW}}$ represent the change in bonded energies (bond, angle, and dihedral), electrostatic energies, and van der Waals energies upon ligand binding, respectively. $\Delta G_{\mathrm{sol}}$ is the sum of the nonpolar solvation energy $\Delta G_{\mathrm{SASA}}$ and electrostatic solvation energy $\Delta G_{\mathrm{GB}}$, computed using the Poisson–Boltzmann method. $\Delta G_{\mathrm{SASA}}$ can be written as-

$$\Delta G_{\mathrm{SASA}} = \gamma SASA + \beta \tag{4}$$

Where $\gamma$ ($= 0.00542$ kcal Å$^2$) is the surface tension, while $\beta = 0.92$ kcal mol$^{-1}$, and SASA represents the solvent-accessible surface area of the molecule. We predicted the binding affinity of DOTMA lipids to the TDN cages from the binding free energy calculation.

Unlike conventional dsDNA, which features two distinct grooves (major and minor) as primary binding sites, a tetrahedral DNA nanostructure, comprising six dsDNA helices and six single-base staples, offers a more complex 3D architecture with multiple accessible binding regions, making it a highly versatile and promising platform for drug delivery. To assess the stability of the binding modes, all-atom molecular dynamics simulations were performed for the TDN–DOTMA complex. We began by examining the interaction of a single DOTMA lipid with TDN, which ultimately favored binding at the outer corner of the nanostructure, with a binding free energy of approximately –32 kcal/mol, indicating a strong binding affinity. This interaction was predominantly driven by electrostatic forces and further stabilized by hydrogen bonds, though partially counteracted by high solvation energy. Overall, electrostatics played the central role in facilitating the association between cationic DOTMA lipids and TDN nucleotides. Detailed energy contributions are summarized in Table St.2.

Table St.2: Different energy components contributing to the average binding free energy of a single DOTMA-TDN complex were calculated over the last 100 ns of the 200 ns MD trajectory at 300 K, expressed in kcal/mol.

| **Component** | **TDN-DOTMA** |
|---|---|
| Van der Waals | $-35.42 \pm 4.04$ |
| Electrostatic | $-1847.05 \pm 40.96$ |
| Generalized Born | $1861.60 \pm 41.12$ |
| $\Delta G_{gas}$ | $-1882.48 \pm 41.70$ |
| $\Delta G_{Solvation}$ | $1857.59 \pm 41.03$ |
| $\Delta G_{Binding}$ | $-24.88 \pm 3.51$ |

We infer that similar binding strengths may be expected for DOTMA lipids in the 10 and 100 DOTMA–TDN systems, despite their involvement in clustered assemblies. Therefore, in the following section, we further analyzed the contributions of non-bonded electrostatic and van der Waals interactions across all complexes, as these are the key components of the binding enthalpy in TDN–DOTMA systems.

Table St.3: Binding Enthalpy energy from the MMGBSA calculation for the three TDN-DOTMA complexes, calculated over the final 50 ns of the 200 ns simulation.

| **System** | **vdW (kcal/mol)** | **ELE (kcal/mol)** | **Enthalpy (kcal/mol)** |
|---|---|---|---|
| **TDN-1 DOTMA** | -35.42±4.04 | -1847.05±40.96 | -1882.48±41.70 |
| **TDN-10 DOTMA** | 742.34±22.42 | -22985.03±220.01 | -22242.68±224.95 |
| **TDN-100 DOTMA** | 2252.05±56.36 | -206371.85±256.19 | -204119.80±277.27 |

### S.7 TDN Complexation with 250 DOTMA Lipids

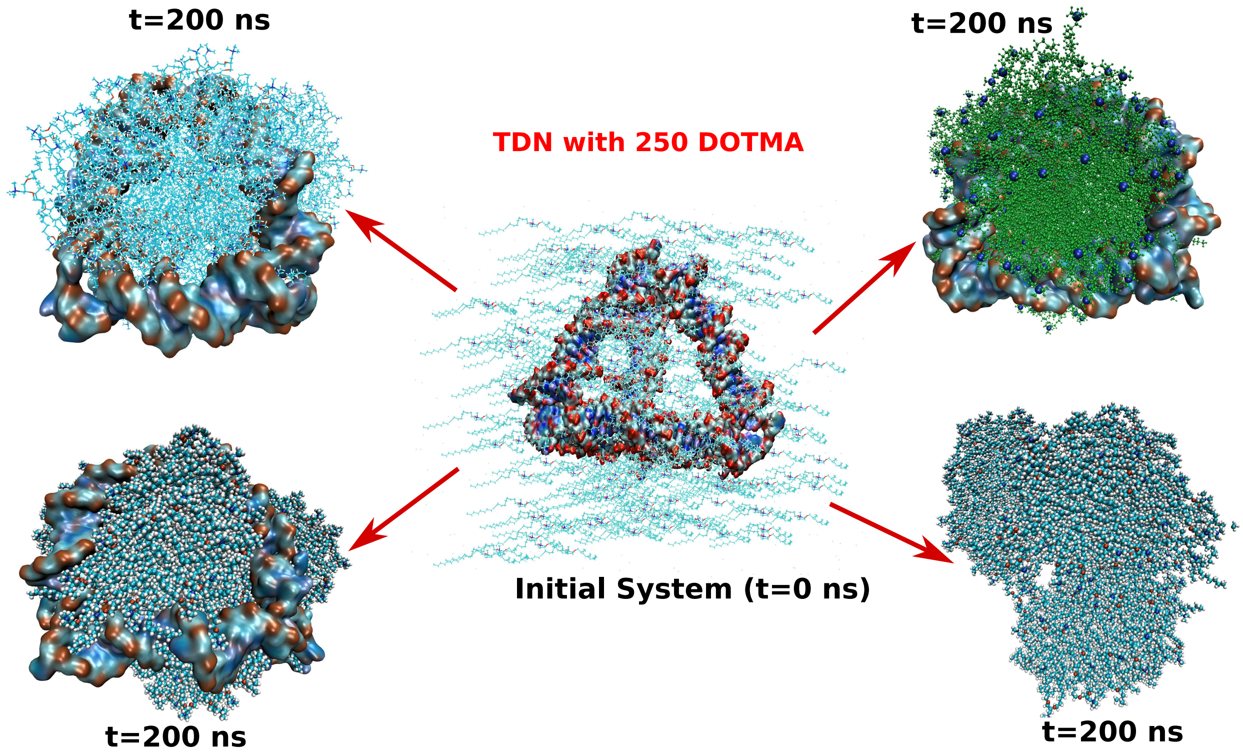


Figure S.5: Initial and Final snapshots of the TDN nanostructures with 250 cationic DOTMA lipid complexes at a physiological salt concentration of 150 mM. The DOTMA lipids maintain their self-assembly and cluster formation and are associated with TDN nucleotides. The lipid cluster is entrapped within the TDN cavity.
